# Bacterial biofilm formation on soil fungi: a widespread ability under controls

**DOI:** 10.1101/130740

**Authors:** Cora Miquel Guennoc, Christophe Rose, Jessy Labbé, Aurélie Deveau

## Abstract

In natural environments, bacteria preferentially live in biofilms that they build on abiotic surfaces but also on living tissues. Although fungi form extensive networks of hyphae within soils and thus could provide immense surfaces for bacteria to build biofilms and to proliferate, the extent on such phenomenon and the consequences for the fitness of both microorganisms is poorly known in soils. Here, we analyzed the process of formation of biofilms by various bacteria on hyphae of soil fungi in an *in vitro* setting using confocal and electron microscopy. We showed that the ability to form biofilms on fungal hyphae is widely shared among soil bacteria. In contrast, some fungi, mainly belonging to the Ascomycete class, did not allow for the formation of bacterial biofilms on their surfaces. The formation of biofilms was also strongly modulated by the presence of tree roots and by the development of the ectomycorrhizal symbiosis, suggesting that biofilm formation does not occur randomly in soil but that it is highly regulated by several biotic factors. Finally, our study led to the unexpected finding that networks of filaments made of extracellular DNA were used to build the skeleton of biofilms by a large array of bacteria.

## Introduction

Among the myriad of organisms that live in forest soils, bacteria and fungi largely exceed their counterparts in terms of abundance and diversity (Nazir *et al*., 2010). Both highly contribute to the decomposition of soil organic matter and to the nutrient cycling and thus have a key role in the modulation of soil fertility and productivity (Rousk and Bengtson, 2014; Lindahl and Tunlid, 2015). In addition, some mutualistic fungi called mycorrhizal fungi, act as providers of carbon sources to the soil and of nutrients to the trees through the symbiosis they establish with roots (Smith and Read, 2008). In soil, many bacteria and fungi often occupy a shared microhabitat and there, many bacteria often colonize the surface of fungal hyphae, also called “hyphosphere” (Frey-Klett *et al*., 2011). Bacteria are thought to gain two main benefits from this association. First, the hyphosphere provide a nutritional source for bacteria that either consume nutrients released directly or indirectly by hyphae, or directly prey on fungi (Leveau and Preston, 2008; Hover *et al*., 2016). Second, fungal hyphae can serve as vectors for bacteria to travel across the soil and to reach otherwise inaccessible nutrient sources (Nazir *et al*., 2010). These so called “hyphal highways”, can be followed by bacteria that swim along the water film that covers the hyphae, or by bacteria that settle at the tip of the growing hyphae (Otto *et al*., 2016; Warmink and van Elsas, 2009). Conversely, some fungi can benefit from the metabolic activity of their associated bacteria (Li *et al*., 2016), gain protection against stresses (Nazir *et al*., 2014) or even “farm” bacteria to later use them as a source of nutrients (Pion *et al*., 2013). However, this close interaction between fungi and bacteria can also be detrimental to the fungi and a number of them produces defensins to prevent the bacterial colonization of their hyphae (Essig *et al*., 2014).

Bacteria can establish in the hyphosphere in three states: as free-living cells, as attached single cells or as organized biofilms. Biofilms arise through the aggregation of bacterial cells and their embedding into a self-produced matrix of extra polymeric substances (Flemming *et al*., 2016). Life as a biofilm has the double advantages to increase the bacterial resistance against biotic and abiotic stresses, and to permit the organization of cells into functional sub-communities. As a consequence, a large number of bacterial species have developed the ability to build biofilms on hydrated abiotic surfaces (e.g. water pipes, medical devices…) but also on living tissues (e.g. epithelial cells, root surfaces…). Fungal hyphae can also support bacterial biofilms and *in vitro* formation of bacterial biofilms on hyphae of some soil Ascomycetes, Basidiomycetes and Zygomycetes has been reported (Scheublin *et al*., 2010; Burmølle *et al*., 2012; Hover *et al*., 2016; Nazir *et al*., 2014). Yet, while soil fungi offer relatively close habitats to bacteria, different behaviors exist between soil fungal species. Since bacterial biofilms are frequent on sporocarps of ectomycorrhizal (ECM) fungi but rare on saprotrophic ones, it has been proposed that the trophic status (i.e. symbiotic vs saprotrophic) of the fungi drives the interactions (de Carvalho *et al*., 2015). As being in competition for carbon sources, saprophyte would be more intolerant to bacteria than mycorrhizal fungi that get their carbon from their plant partner. However, this hypothesis has not been tested on hyphae yet. Conversely, differences among bacteria in their ability to form biofilms on hyphae have been pointed in the arbuscular mycorrhizal (AM) fungi *Rhizophagus intraradices* and *Glomus* sp. (Toljander *et al*., 2006; Scheublin *et al*., 2010) suggesting a potential fine-tuned interaction between these fungi and bacteria during the process of biofilm formation. Specific bacterial communities also preferentially associate with ECM fungi (Frey-Klett *et al*., 2005; Warmink *et al*., 2009). Among the ECM associated bacteria, Mycorrhiza Helper Bacteria (MHB) have received a lot of attention because of their ability to stimulate mycorrhiza formation and functioning (Deveau and Labbé, 2017). Some of them have been shown to form biofilm-like structure on ECM fungi although the extent of such interactions is unknown (Frey-Klett *et al*., 2007). Because ECM fungi colonize very large volumes of soil, they could provide large surfaces for bacteria to establish as biofilms. Altogether, these data suggest that biofilm formation on fungal hyphae could be an important phenomenon in soil. However, we have little information on the frequency of such event, its level of specificity among microorganisms and the biotic and abiotic parameters that drive the fate of the interaction. Such knowledge is key to predict how soil fungi and bacteria behave in soils.

To fill this gap, we characterized in depth the biofilms formed by several MHBs and other soil bacteria on hyphae of the model ECM fungus *Laccaria bicolor* S238N (Martin *et al*., 2008) using an *in vitro* set up (Miquel Guennoc *et al* 2016). We then evaluated the impact of the root system and of the ectomycorrhizal symbiosis on the fate of the interaction. Lastly, we tested the impact of the trophic status and the taxonomic origin of soil fungi on the establishment of biofilms. Our data suggest that biofilm formation on hyphae does not occur randomly but then it is regulated by several biotic factors. In addition, the detailed study of the structure of the bacterial biofilms revealed an unexpected feature that is likely to be widespread: the use of extracellular DNA filaments to build the skeleton of biofilms.

## Material and methods

### Microbial strains and culture conditions

All the strains used in this study are listed in Table 1. GFP-tagged versions of *Pseudomonas fluorescens* BBc6 (Deveau *et al*., 2010), *Burkholderia ginsengisoli* E3BF7_7 and Dyella sp. E3BF9_7 were used to facilitate imaging. None of the other microbial strains used in this study constitutively expressed a fluorescent protein. Bacterial strains were maintained at -80°C in Luria-Bertani (LB) medium with 30% glycerol and were first grown on 10% tryptic soy agar (TSA)-plates for 24h (3 g.l^-1^ tryptic soy broth from Difco and 15 g.l^-1^ agar) at 28°C. Then, for each strain, 2 to 3 individual bacterial colonies were collected from TSA cultures to inoculate 25 ml of liquid LB medium and incubated at 28°C and 150 rpm until late exponential growth before their use for biofilms formation. Fungal cultures were maintained on P5 medium then transferred to P20 agar plates covered with EDTA pre-treated cellophane membranes as described by Miquel Guennoc *et al*. (2016), except for *Tuber melanosporum* that was always kept on P20 medium.

**Table 1.** List and characteristics of the microbial strains.

| Microbial strains | Taxonomic classification | Gram type | Ecological trait | References |
| --- | --- | --- | --- | --- |
| **Bacterial strains** |  |  |  |  |
| *Pseudomonas fluorescens* BBc6 | Gamma-proteobacteria, Pseudomonadaceae | - | Mycorrhiza helper bacterium | Duponnois & Garbaye 1991 |
| *Pseudomonas protegens* Pf5 | Gamma-proteobacteria, Pseudomonadaceae | - | Biocontrol | Howell & Stipanovic 1979 |
| *Pseudomonas fluorescens* SBW25 | Gamma-proteobacteria, Pseudomonadaceae | - | Plant Growth Promoting & biocontrol | Bailey *et al* 1995 |
| *Pseudomonas fluorescens* Pf29A | Gamma-proteobacteria, Pseudomonadaceae | - | Biocontrol | Chapon *et al* 2002 |
| *Pseudomonas* sp. GM18 | Gamma-proteobacteria, Pseudomonadaceae | - | Mycorrhiza helper bacterium | Labbe *et al* 2014 |
| *Pseudomonas syringae* pv. *tomato* DC3000 | Gamma-proteobacteria, Pseudomonadaceae | - | Plant pathogen | Cuppels 1986 |
| *Dyella* sp. E3BF9_7 | Gamma-proteobacteria, Rhodobacteraceae | - | White rot associated bacterium | Hervé *et al* 2016 |
| *Collimonas fungivorans* Ter331 | Beta-proteobacteria, Oxalobacteraceae | - | Mycophagous bacterium | de Boer *et al* 2004 |
| *Burkholderia ginsengisoli* E3BF7_7 | Beta-proteobacteria, Burkholderiaceae | - | White rot associated bacterium | Hervé *et al* 2016 |
| *Sinorhizobium meliloti* 1021 | Alpha-proteobacteria, Rhizobiaceae | - | Nitrogen fixing plant symbiont | Maede *et al* 1982 |
| *Pedobacter* sp. D3AIN17A | Bacteroidetes, Sphingobacteriaceae | - | Black truffle associated bacterium | Deveau *et al* unpublished |
| *Paenibacillus* sp. F2001-L | Firmicutes, Paenibacillaceae | + | Endohyphal strain of *L. bicolor* | Bertaux *et al* 2003 |
| *Bacillus subtilis* MB3 | Firmicutes, Bacillaceae | + | Mycorrhiza helper bacterium | Duponnois & Garbaye 1991 |
| *Bacillus* sp. EJP109 | Firmicutes, Bacillaceae | + | Mycorrhiza helper bacterium | Poole *et al* 2001 |
| **Fungal strains** |  |  |  |  |
| *Laccaria bicolor* S238N | Basidiomycota, Tricholomataceae | n/a | ECM | Martin *et al* 2008 |
| *Lactarius quietus* | Basidiomycota, Russulaceae | n/a | ECM | INRA Nancy |
| *Hebeloma cylindrosporum* | Basidiomycota, Cortinariaceae | n/a | ECM | Kohler *et al* 2015 |
| *Piloderma croceum* | Basidiomycota, Atheliaceae | n/a | ECM | Kohler *et al* 2015 |
| *Phanerochaete chrysosporium* RP78 | Basidiomycota, Phanerochaeteaceae | n/a | saprotroph (white rot) | Hervé *et al* 2016 |
| *Thelephora terrestris* | Basidiomycota, Thelephoraceae | n/a | ECM | INRA Nancy |
| *Tuber melanosporum* Mel28 | Ascomycota, Tuberaceae | n/a | ECM | Martin *et al* 2010 |
| *Elaphomyces granulatus* PEP | Ascomycota, Elaphomycetaceae | n/a | ECM | Quandt *et al* 2015 |
| *Penicillium funiculosum* PF01 | Ascomycota, Aspergillaceae | n/a | Saprotroph & plant pathogen | INRA Nancy |
| *Aspergillus ustus* AU01 | Ascomycota, Aspergillaceae | n/a | Saprotroph | INRA Nancy |

### In vitro biofilm formation on fungal hyphae and glass fibers

The method described step-by-step in Miquel Guennoc *et al*. (2016) was used. Briefly, bacterial cultures in late exponential growth phases were spin down and washed once in potassium phosphate buffer (PPB; KH_2_PO_4_ 25 g.l^-1^, K_2_HPO_4_ 2.78 g.l^-1^, pH 5.8). Bacterial pellets were suspended in PPB and the cell density was adjusted to 10^9^ cfu.ml^-1^ to prepare the bacterial inoculum. Five ml of this bacterial suspension was added to each well of 6 well-microplates. Then, using sterile tweezers, 1-cm diameter fungal colonies grown on P20-agar plates were added to each well of the microplates. The cellophane membranes on which they were growing were kept to avoid harming fungal hyphae during the transfer. Plates were gently shaked for 1 min to allow the fungus to unstick from the cellophane sheets and cellophane sheets were removed from each well. These resulting microplates with the bacterial and fungal inocula were incubated at 20°C and 60 rpm for 16hrs, except if otherwise stated. To test for biofilm formation on dead fungal colonies, 1-cm diameter growing fungal colonies were dipped into either 3% paraformaldehyde solution for an hour or progressive ethanol baths for 3 min each (20%, 50%, 70%, 100 %), then washed three times with PPB to remove traces of paraformaldehyde or ethanol,. Dead colonies were then immediately used to test for biofilm formation using same protocol as above. To test the formation of bacterial biofilm on glass fibers, fungal colonies were replaced by 10-μm diameter sterile glass fibers. For each treatment, two independent assays in triplicate were analyzed.

### Sample preparation for confocal and electron microscopy imaging

Samples for imaging were prepared following the methodology described in Miquel Guennoc *et al*. (2016). Briefly, fungal colonies or glass fibers were transferred to a new microplate. Then, samples were rinsed with NaCl (17 g.l^-1^) and with PPB to detach planktonic and electrostatically attached bacterial cells (Toljander *et al*., 2006).

For confocal imaging, fungal colonies were cut in half with a razor blade, stained then mounted on slide with Fluoromount-G anti-fading (Fisher Scientific). Fungal hyphae were stained with 10 μg.ml^-1^ Wheat Germ Agglutinin coupled to Alexa Fluor 633 (WGA-633, Thermofisher Scientific) for 15 min. All bacteria, except *P. fluorescens* BBc6, *B. ginsengisoli* E3BF7_7 and Dyella sp. E3BF9_7 that constitutively expressed GFP, were stained with 0.3 μM 4’,6-Diamidino-2-Phenylindole (DAPI, Thermofisher Scientific) for 15 min. Several dyes were used to visualize matrix components: 1X SYPRO Ruby (15 min incubation; Thermofisher Scientific), DAPI, prodidium iodide (1μg.ml^-1^, 15 mins; Thermofisher Scientific) or TO-PRO-3 (1 μM, 15 mins; Thermofisher Scientific) were used to stain proteins and extracellular DNA (eDNA), respectively (Suppl. Table 1). Confocal imaging was performed with a LSM780 Axio Observer Z1 laser scanning confocal microscope (LSCM, CarlZeiss), equipped with 405, 488 and 633 nm excitation lasers and T-PMT and GaAsp PMT detectors, coupled to ZEN 2.1 lite black software (CarlZeiss). For all experiments, images were captured with 10x 0.3 NA objective to obtain a complete view of one fourth of the fungal using a combination of tile scan and Z stack functions (5x5 fields over the entire depth of the fungal colony). Then images were taken with a 40x 1.2 NA objective to obtain high resolution zoom images of representative events within the imaged captured at 10x. Data visualization was performed by 2D maximum intensity projection, using the “Z project” function from Fiji free software ((Schindelin *et al*., 2012), http://fiji.sc/Fiji).

For electron microscopy imaging, fungal colonies or glass fibers were first rinsed with NaCl (17 g.l^-1^) and with sterile water to avoid crystal formation during dehydration step. Then, samples were dehydrated by freeze-drying and coated with 2 nm of platinum (quartz measurement) under argon plasma (2.5.10-2 mbar, 35 mA) with a High Vacuum Coater Leica EM ACE600 (Leica). Coated samples were imaged first with a scanning electron microscope (SEM) equipped with a Field Emission Gun (SIGMA-HPSEM-FEG, Zeiss) using high resolution “in lens” detector at 1 kV of accelerating voltage. In a second step, some samples (biofilms grown on glass fibers) were placed in a second SEM (LEO 1450VP W-SEM, Zeiss) to perform EDS micro-analysis and mappings of elements (20 kV of accelerating voltage at 1 nA of sample current; Oxford-Instruments INCA MAPS software)

### Biofilm formation in the presence of Populus roots and ectomycorrhizae

Micro-propagated hybrid poplar (*Populus tremula* × *Populus alba*; INRA clone 717-1-B4) were used to form mycorrhiza with *L. bicolor* S238N following the method of “*in vitro* sandwich co-culture system” developed by Felten *et al*. (2009). A mycelium-covered cellophane membrane was placed fungus side down on the roots of 3 weeks old *Populus* seedlings. Petri dishes were closed with Band-Aids (ensuring high gas permeability). Cultures were arranged vertically, and the lower part of the dish was covered with a small black plastic bag to prevent light from reaching the fungus and roots. The co-cultures were incubated for one month at 24°C and under 16hrs-,photoperiod then the development of mature ectomycorrhizae was controlled under a stereoscope (CarlZeiss). At this stage, bacterial suspensions were prepared as described above. Seedlings of *Populus* colonized by *L. bicolor* were transferred in a large Petri dish filled with sterile PPB (Suppl. Fig. 1). The cellophane membranes were gently detached by agitation (1 min) and removed. Mycorrhizal seedlings were transferred into a double Petri dish setting containing the bacterial suspension (Suppl. Fig. 1). The double Petri dish was designed to prevent contact between plant shoot and bacterial suspension. The systems were then incubated at 20°C with gentle agitation (60 rpm) for 16h. Control treatments (*Populus* plants not inoculated with *L. bicolor*) were treated similarly. After 16h of incubation, mycorrhizal and non-mycorrhizal seedlings were transferred in a new double Petri dish to be washed with NaCl and PPB as described above. Each sample was then examined by confocal microscopy. To obtain transversal cross sections of ectomycorrhizae, ectomycorrhizae were included in 4% agarose and sectioned with a vibratome (Leica VT1200S) at a thickness of 30 μm. For each treatment, two independent assays in duplicate were performed.

### Enzymatic treatment of biofilms

To investigate the role of extracellular DNA (eDNA) in biofilms formation, DNase I (30 Kunitz units.ml^-1^ Qiagen) was added to *P. fluorescens* BBc6 suspensions before the incubation with glass fibers. To analyze the composition of *P. fluorescens* BBc6 biofilm filaments, biofilms grown for 16 hrs on glass fibers were treated with proteinase K (60 mAnson units.ml^-1^ for 1 hr, ThermoFisher), DNase I (30 Kunitz units.ml^-1^ Qiagen), RNase A (100 Kunitz units.ml^-1^, ThermoFisher) or cellulase R10 (1 mg.ml^-1^, from *Trichoderma viride*, SERVA). All the enzymatic treatments were performed at room temperature for 2hrs. Samples were observed with laser scanning confocal microscopy (LSCM). Positive controls were treated similarly. Proteinase K activity was verified with non-fat dried skimmed milk powder as described by Nygren *et al*. (2007). Cellulase activity was verified with AZCL-HE-Cellulose per the manufacturer’s instructions (0.2 % w/v, Megazyme). For RNase activity, total RNA from human placenta (1 μg.μl^-1^, Clontech) was treated and degradation was verified by electrophoretic migration of treated and non-treated RNA (negative control) in 1 % -agarose gel stained with ethidium bromide.

## Results

### Biofilm formation on L. bicolor S238N hyphae is widespread among soil bacteria

We previously reported that the MHB *P. fluorescens* BBc6 forms biofilm structures on the hyphae of *L. bicolor* S238N in *in vitro* setting (Miquel Guennoc *et al*., 2016). Using the same setting, we further characterized the process by Laser Scanning Confocal Microscopy (LSCM) and Scanning Electron Microscopy (SEM). The attachment of individual bacterial cells to the surface of hyphae was observed after a few minutes of contact between the microorganisms (Fig. 1a, d), followed by the formation of colonies (after few hours) made of a matrix of several layers of cells engulfing the hyphae (Fig. 1b, e). The colonies kept building up to form mature biofilms after about 20 hours (Fig. 1c, f). Bacterial cells were encased in a dense matrix of extracellular polymeric substances (Fig. 2a) made of extracellular DNA (eDNA) and proteins (Fig. 2b). Filaments that bound DAPI stain connected bacterial cells to each other and anchored the biofilms to the surfaces of the hyphae (Fig. 2 a, c). The substitution of *L. bicolor* hyphae by glass fibers (similar diameter) in controls also led to biofilm formation around the glass fibers by *P. fluorescens* BBc6 (Fig. 3a), potentially suggesting that fungal hyphae may be nothing more than a physical support used by bacteria to establish biofilms. To further test this hypothesis, biofilm formation on alive and dead fungal colonies was compared. *P. fluorescens* BBc6 formed biofilms on both living and dead hyphae but the distribution of the biofilms greatly differed between the two treatments. Although sparse attachment was detected all over the fungal living colonies, mature biofilms only developed at the actively growing margin of the fungal colony (Fig. 3b). In contrast, biofilms were found all over dead fungal colonies (Fig 3c) suggesting that specific interactions between bacterial and fungal cells occur during the formation of biofilms and that fungal hyphae are more than physical supports.

**Figure 1.**
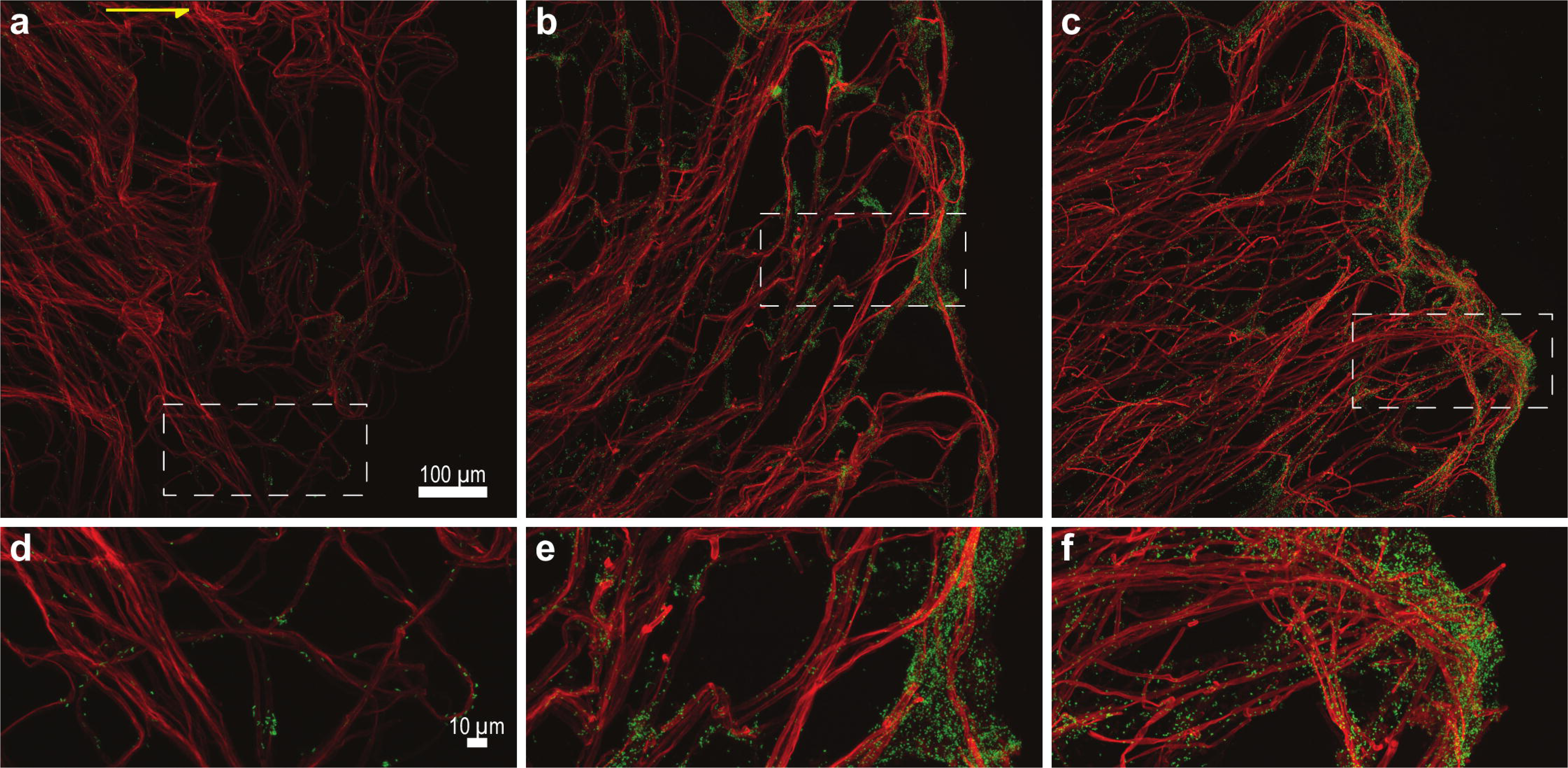
Time course of *P. fluorescens* BBc6 biofilm formation on the surface of *L. bicolor* S238N hyphae. Confocal microscopy images showing the spatial localization of BBc6 biofilms (green) on *L. bicolor* S238N hyphae (red) and their development over time from early stage attachment (30 min; a,d) to colony formation (6h; b,e) and mature biofilms (20h; c,f). The yellow arrow points toward the external edge of the fungal colony. Bottom panels are zoom in of the areas highlighted by white rectangles in top panels. Fungal hyphae were stained with Wheat Germ Agglutin-AlexaFluor 633 and bacterial cells were GFP-tagged. Images were obtained via 2D maximum intensity projection of 3D confocal microscopy images (z = 40.5μm (a), 30μm (b), 43.5μm (c)). Magnification 40x.

**Figure 2.**
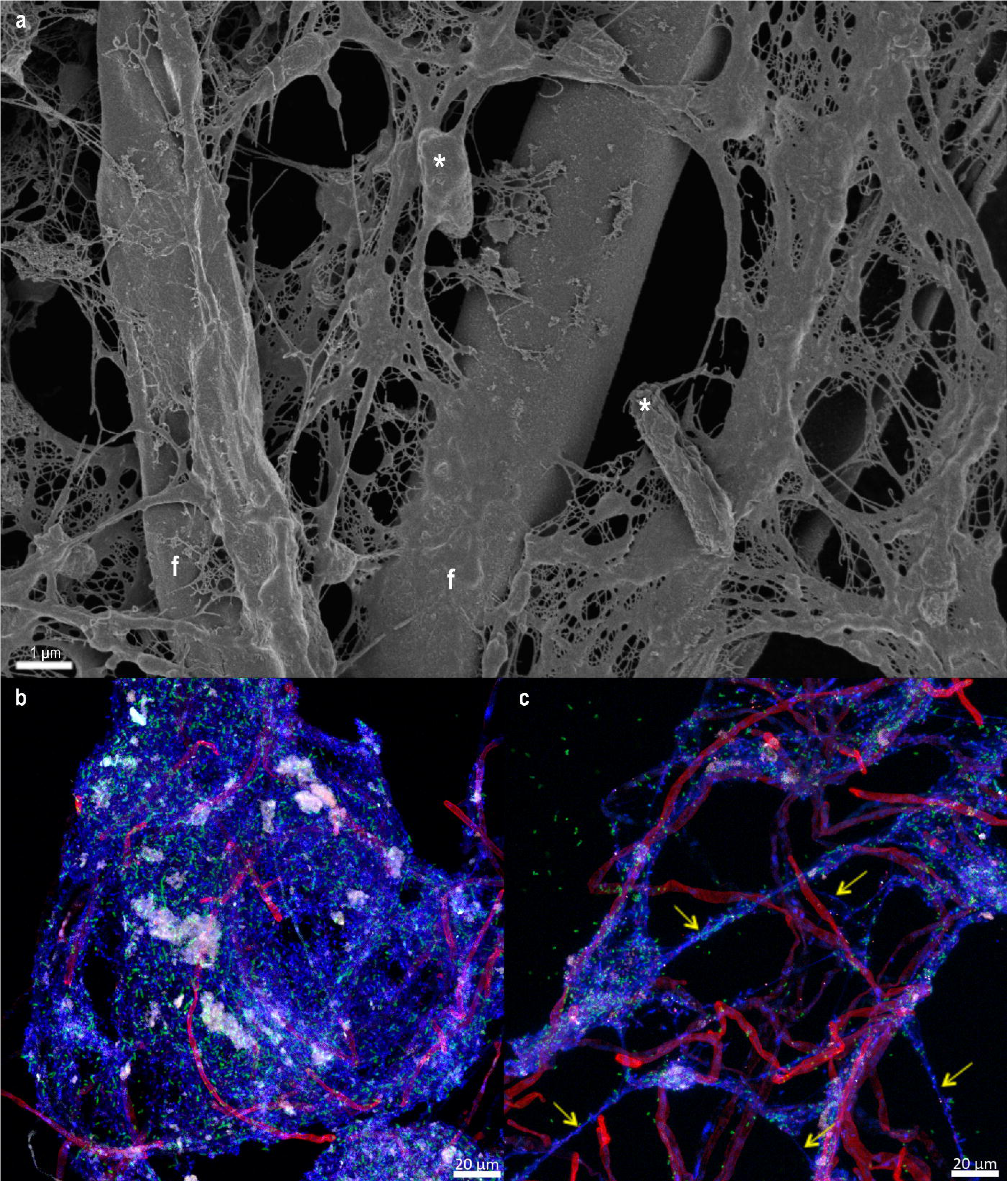
Characterization of the matrix components of *P. fluorescens* BBc6 biofilms on the surface of *L. bicolor* S238N. **A.** Scanning electron microscopy image of 24h old biofilm showing bacterial cells (*) and fungal hyphae (f) encased in a matrix made of aggregates and filaments. **B.** Confocal microscopy image showing the presence of proteins aggregates (white) and eDNA (blue) in the matrix of 16h old *P. fluorescens* BBc6 (green) biofilm on *L. bicolor* S238N hyphae (red). Proteins and eDNA were stained with SYPRO Ruby and DAPI, respectively. Magnification 40x. **C.** Confocal microscopy image showing the presence of filaments stained by DAPI (yellow arrows) connecting hyphae to bacterial cells. Magnification 40x.

**Figure 3.**
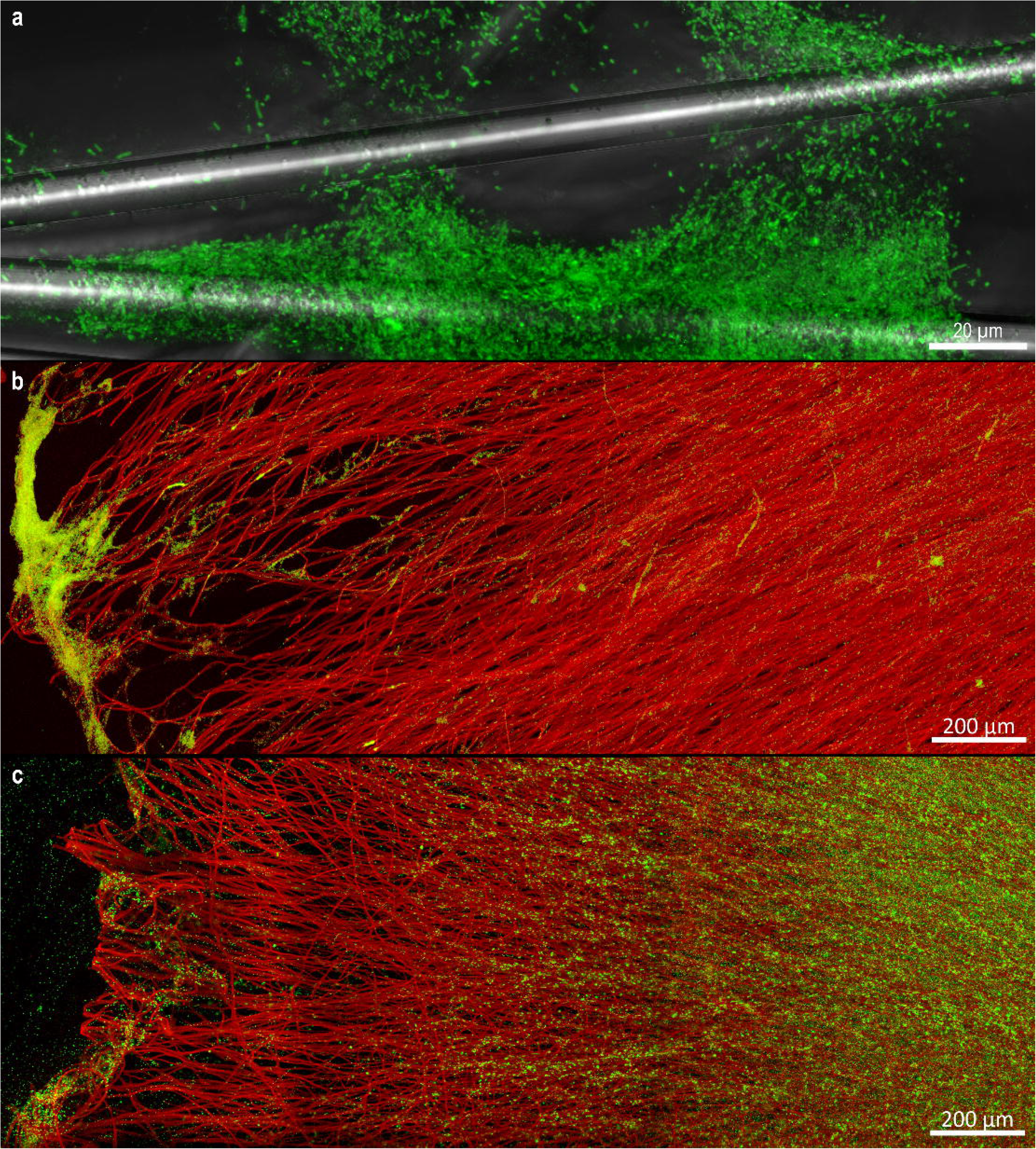
Distribution of *P. fluorescens* BBc6 biofilms on abiotic and biotic surfaces. **A.** Confocal microscopy image showing *P. fluorescens* BBc6 biofilm formed over glass fibers after 22 hours. Magnification 40x. **B, C.** Confocal microscopy images showing differential distribution *P. fluorescens* BBc6 biofilms formed over alive (B) and dead hyphae (C) of *L. bicolor* S238N. *L. bicolor* S238N hyphae were killed by immersing fungal colonies in 3% paraformaldehyde for 1h followed by 3 repeated washes in phosphate buffer before inoculating bacteria. Images were obtained via 2D maximum intensity projection of 3D mosaic confocal microscopy images (z = 69μm (b), 32 μm (c)). Magnification 10x.

To assess the degree of specificity of the physical interaction between the MHB and the ECM fungus, thirteen additional cultivable bacteria spanning over a wide range of taxa highly represented in soil or associated with plants, and with various ecological traits (e.g. MHB, biocontrol, pathogen, Table 1) were tested for their ability to form a biofilm on *L. bicolor* S238N hyphae. All bacteria formed biofilms around the hyphae at the edge of *L. bicolor* colonies (Suppl. Fig 2).

### Skeletons of bacterial biofilms are made of DNA filaments

Biofilms formed by the 14 bacterial strains on hyphae of *L. bicolor* S238N and on glass fibers were all characterized by the presence of complex networks of filaments stained by DAPI (Suppl. Fig 2). These filaments could reach a length of several hundred micrometers, and were produced, at least, by the bacteria since they were also retrieved in BBc6 biofilms formed on glass fibers (Fig. 4a). The filaments could only be visualized when stained with DNA specific dyes (DAPI, propidium iodide and TO-PRO-3, Suppl. Table 1A). None of the other dyes tested (e.g. cellulose specific dyes) gave a positive result (Suppl. Table 1A). In accordance to a DNA composition of the filaments, phosphorous and nitrogen, two major components of DNA, were both detected at the surface of the filaments by EDS-SEM elemental mapping (Suppl. Fig. 3). Lastly, a DNAse treatment dismantled the biofilms and disrupted the filaments (Fig. 4b, c) while proteinase K, RNAse and cellulase had no visible effect (Suppl. Table 1B). The filaments were also detected when 2% glucose was added to the incubation medium (data not shown), indicating that the DNA filaments were not produced as a substitution strategy for cellulose caused by the absence of carbon sources (Serra *et al*., 2013).

**Figure 4.**
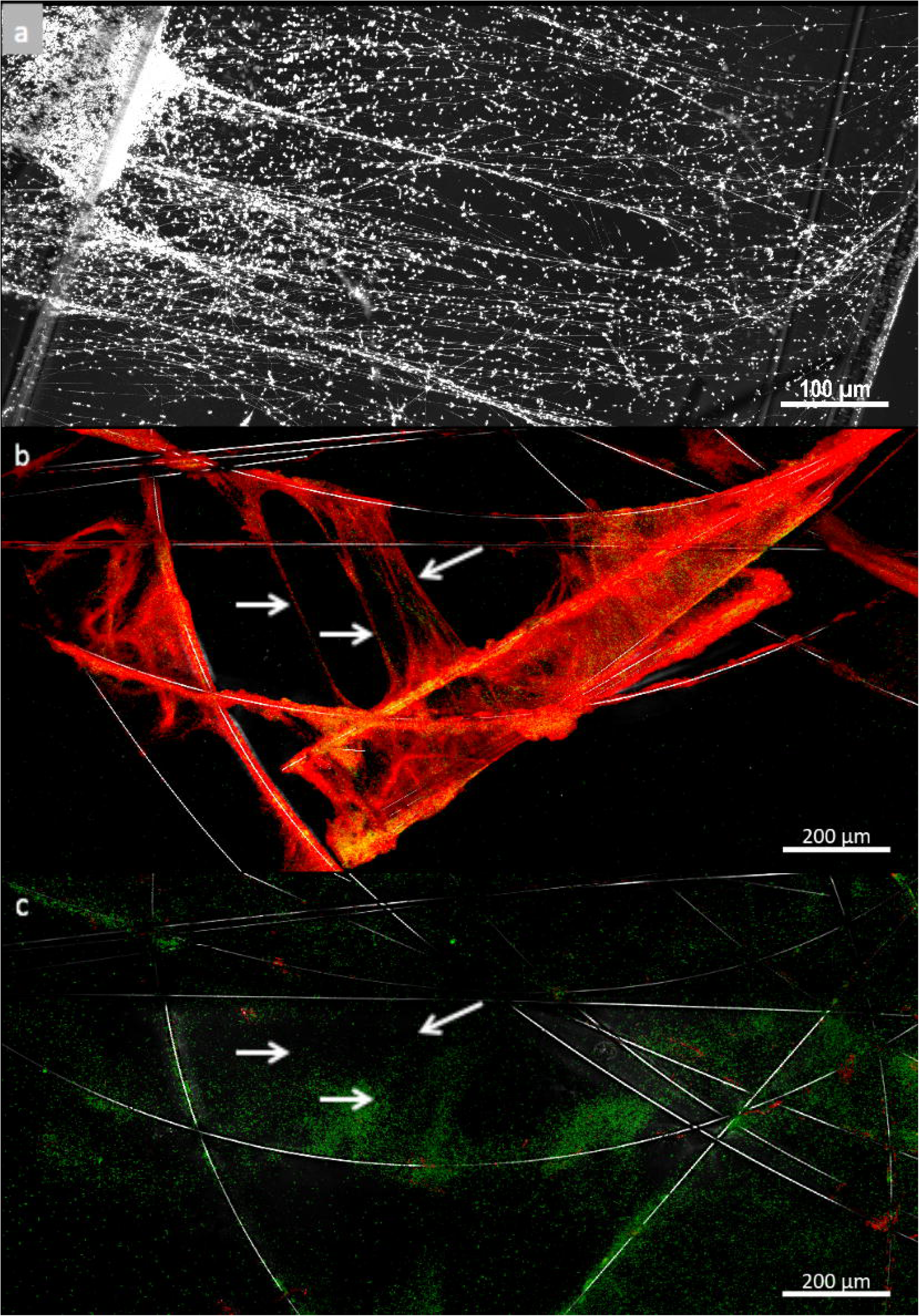
Production of eDNA filaments by *P. fluorescens* BBc6 during biofilm formation. **A**. Confocal images showing mm long filament structures stained by DAPI DNA marker in biofilms of *P. fluorescens* built on glass fibers. Magnification 40x. **B, C.** Confocal images of TO-PRO-3 stained filaments in *P. fluorescens* BBc6 16 hrs old biofilm on glass fibers before (B) and after DNAse treatment (C). White arrows point at filament positions. Magnification 10 x.

### Bacterial biofilm formation on hyphae is restricted to some fungi

We next tested how widespread is the formation of bacterial biofilm on the hyphae of ten ECM and non-ECM soil fungi using *P. fluorescens* BBc6 as a model bacterial strain (Table 1). The bacterial strain formed biofilms on the hyphae of all ECM strains (Table 2, Suppl. Fig. 4) except the Ascomycete *Tuber melanosporum* (Fig. 5a). Conversely, the bacterial strain produced biofilms on the surface of the Basidiomycete wood decay *Phanerochaete chrysosporium* but not on the hyphae of the Ascomycete saprophytes *Aspergillus ustus* AU01 and *Penicillium funiculosum* PF01. In these latter cases *P. fluorescens* BBc6 only attached to the hyphae and never built multilayer biofilms (Fig. 5b). This contrasts with Tuber hyphae on which no attachment at all was visible (Fig 5a).

**Table 2.** Taxonomy and trophic status of fungi tested for bacterial (*P. fluorescens* BBc6) biofilm formation.

| Trophic status & taxonomy | Total of strains tested | Biofilm positive | Biofilm negative |
| --- | --- | --- | --- |
| ECM | 7 | 6 | 1 |
| Ascomycota | 2 | 1 | 1 |
| Basidiomycota | 5 | 5 | 0 |
| Saprotroph | 3 | 1 | 2 |
| Ascomycota | 2 | 0 | 2 |
| Basidiomycota | 1 | 1 | 0 |
| Taxonomy |  |  |  |
| Ascomycota | 4 | 1 | 3 |
| Basidiomycota | 6 | 6 | 0 |

**Figure 5.**
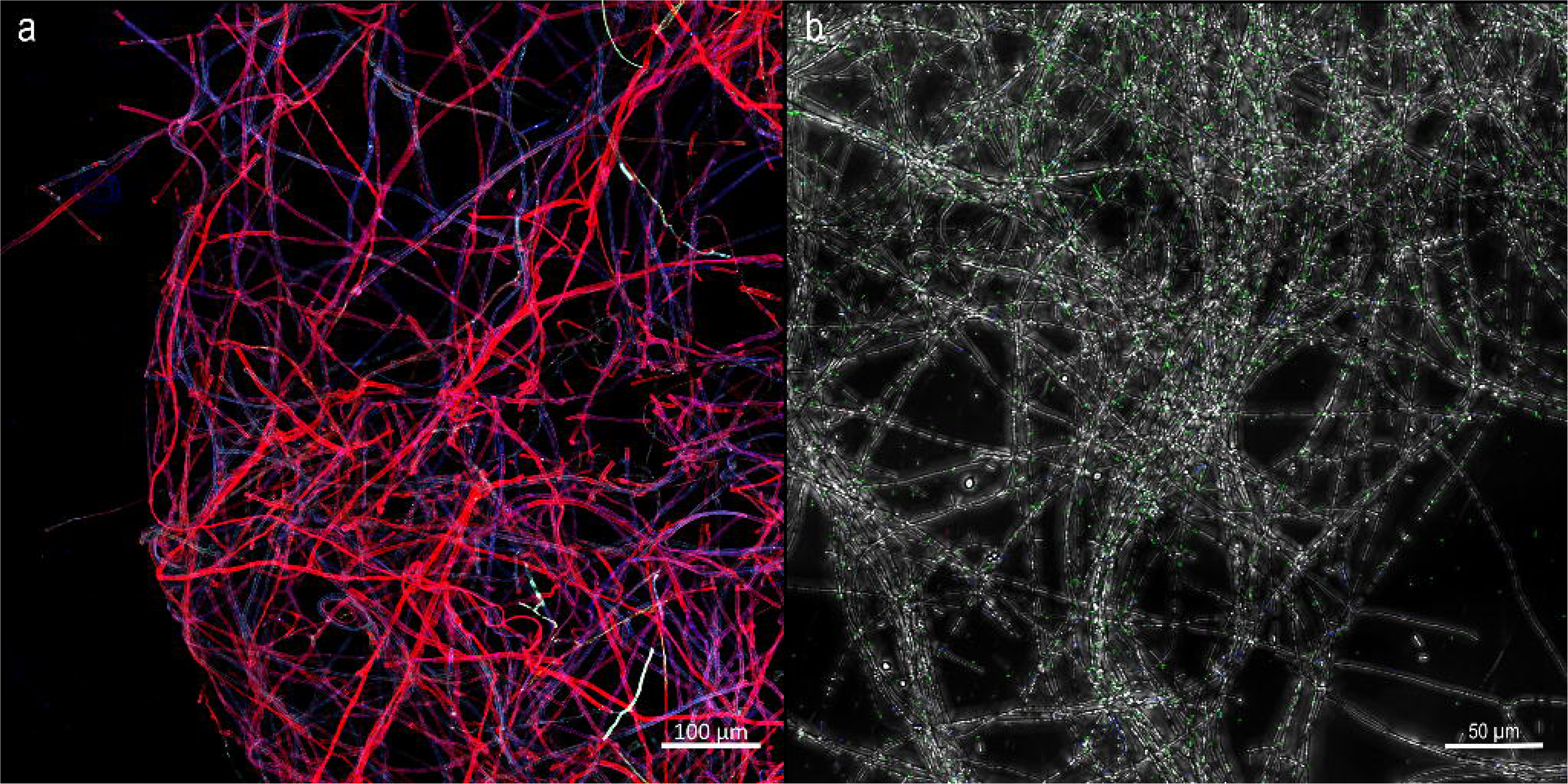
*P. fluorescens* BBc6 does not form biofilm on the hyphae of the Ascomycetes *Tuber melanosporum* (A) and *Aspergillus ustus* (B) after 16hrs of interaction. *T. melanosporum* hyphae were stained with Wheat Germ Agglutin-AlexaFluor 633 (red). Due to poor staining of *A. ustus* hyphae by Wheat Germ Agglutin-AlexaFluor 633, A. ustus hyphae were imaged using transmitted light. Magnification 40x.

### The presence of tree roots and ectomycorrhizae modifies P. fluorescens BBc6 behavior

Roots and ectomycorrhizae are nutrient hotspots that chemoattract complex communities of bacteria and that can provide to bacteria alternative habitats from the hyphosphere (Danhorn and Fuqua, 2007; Bonfante and Anca, 2009). We tested whether the presence of these organs would modify the behavior of *P. fluorescens* BBc6 using Poplar as a tree model organism. Poplar seedlings were grown *in vitro* and used to produce ectomycorrhizae with *L. bicolor* S238N. The root system, together with the associated mycelium, were then incubated with *P. fluorescens* BBc6 bacteria in a liquid setup without nutrients for 16 hours to assess biofilm formation on free roots, ectomycorrhizae, short- and long-distance extramatrical mycelium (Suppl. Fig. 1). Bacteria heavily colonized the surfaces of free roots (Fig 6a), ectomycorrhizae (Fig. 6b), and short-distance extramatrical mycelium (i.e. emerging from the ECM; Fig. 6c). By contrast, no biofilm was detected on distant hyphae (Fig 6d). To test whether this absence of biofilm on distant hyphae was caused by a difference of physiology between short- and long-distance extramatrical hyphae, part of the long-distance extramatrical mycelium was sampled and transferred to a new plate containing a bacterial inoculum. After 16 hours, a biofilm had formed on the surface of this “free” long-distance extramatrical mycelium (Fig. 6e), suggesting that the presence of ectomycorrhizae but not a change in the physiological state of the fungus was responsible for the change of behavior of the bacterium.

**Figure 6.**
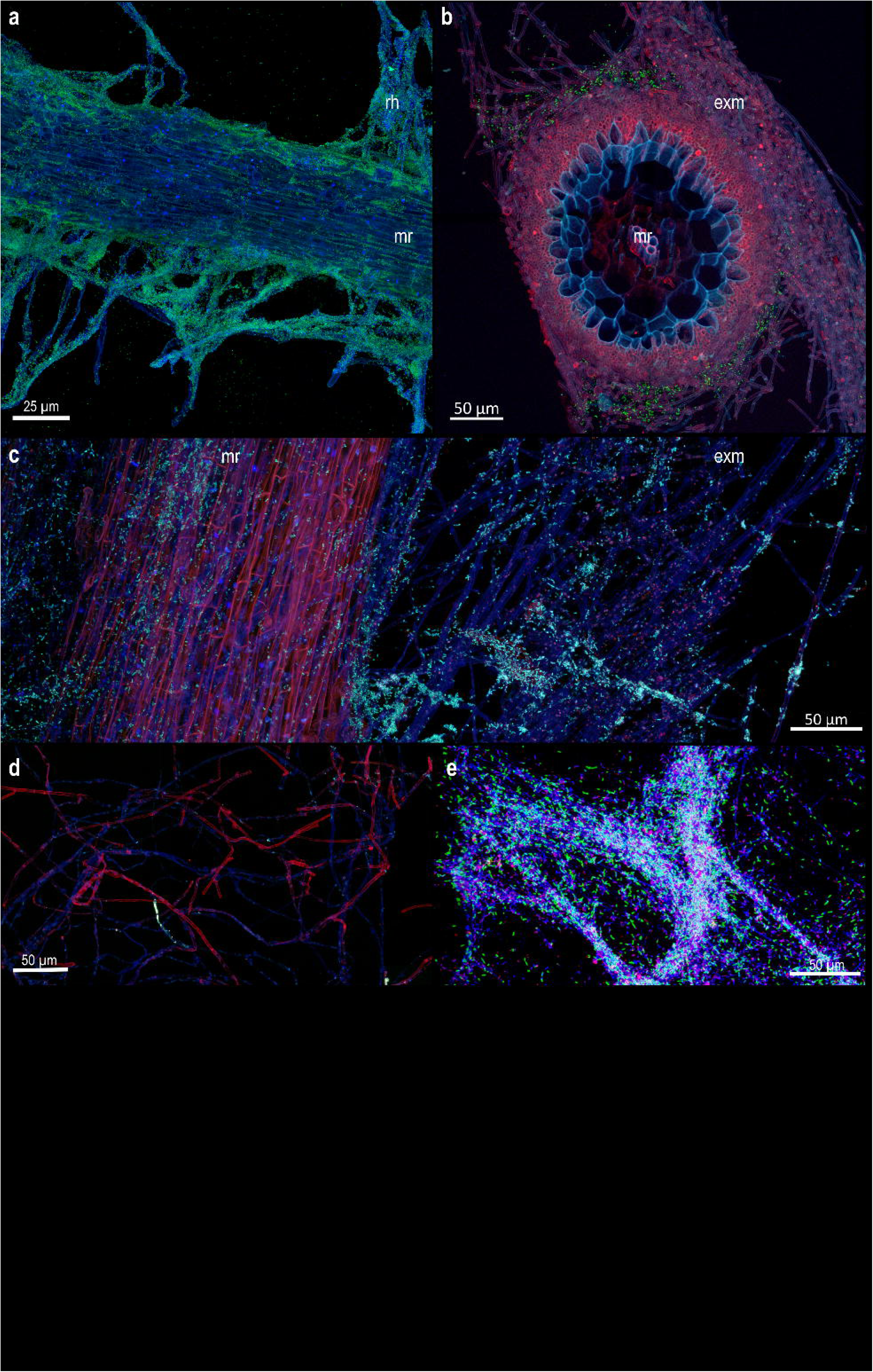
Differential distribution of *P. fluorescens* BBc6 biofilms on Poplar roots, ectomycorhizae and extramatrical mycelium after 16 hrs of interaction. **A.** Confocal image showing *P. fluorescens* BBc6 (green) colonization of Poplar roots (blue). **B.** Transversal section of *L. bicolor* S238N (red)– Poplar (blue) ectomycorrhizae colonized by *P. fluorescens* BBc6 (green). **C.** Confocal image of *L. bicolor* S238N extramatrical hyphae (blue) surrounding Poplar root (red) and colonized by *P. fluorescens* BBc6 (green). **D.** Confocal image showing *L. bicolor* S238N extramatrical hyphae (red) distant from root and ectomycorhizae that are not colonized by *P. fluorescens* BBc6 (green) in the presence of Poplar root system. **E.** Confocal image showing *L. bicolor* S238N extramatrical hyphae (red) distant from root and ectomycorhizae colonized by *P. fluorescens* BBc6 (green) in the absence of Poplar root system. Magnification 40x. *L. bicolor* S238N hyphae were stained with Wheat Germ Agglutin-AlexaFluor 633 (red), Poplar root cells and eDNA were visualized with a combination of DAPI staining and autofluorescence (blue) and bacterial cells were GFP-tagged (green). All images were obtained via 2D maximum intensity projection of 3D confocal microscopy images (z = 60 μm (a), 57 μm (b), 68 μm (c), 46 μm (d), 29 μm (e)). Magnification 40x. mr: main root, rh: root hairs, exm : extramatrical mycelium.

## Discussion & Perspectives

In various natural environments, bacteria preferentially live in biofilms that are built on abiotic surfaces but also on living tissues such as roots (Flemming *et al*., 2016; Burmølle *et al*., 2012). Filamentous fungi represent up to 75% of the subsurface microbial biomass with extended networks of 10^2^ to 10^4^ m length per g of topsoil (Ritz and Young, 2004) and thus could provide immense surfaces for bacteria to form biofilms. Yet, little is known on the extent on such phenomenon and the consequences for the fitness of both microorganisms. Our results confirm that life as a biofilm is imprinted in the genomes of a large taxonomic range of soil bacteria with various life styles and that they are likely to use the surface of hyphae of fungi such as *L. bicolor* to build biofilms. Our data indicate that such biofilm formation is likely to occur on both living and dead hyphae (Fig 3). This contrasts with previous reports on the behavior of *Salmonella enterica* and *Bacillus subtilis* that only formed biofilms on living hyphae of *Aspergillus niger* (Balbontín *et al*., 2014; Benoit *et al*., 2015). However, the use of heat treatment to kill A. niger may have caused a bias by denaturing the structure of the fungal cell wall and thus preventing the molecular interaction between bacterial cellulose and fungal chitin (Balbontín *et al*., 2014). In the present study, bacteria preferentially formed biofilms at the edge of the actively growing colonies while they colonized the entire fungal colonies when those were killed by fixation in paraformaldehyde (Fig 3) or ethanol (data not shown). Formation of biofilms on the growing tips of hyphae of fungi by the bacterium *Burkholderia terrae* was also previously reported (Nazir *et al*., 2014). Such differential distribution could be caused by a polarized heterogeneity of the fungal colony, either in the composition of the fungal cell wall (Latgé, 2007) or in the secretion of nutrients (Webster and Weber, 2007). The localization of bacterial biofilms on fungal hyphae was also strongly influenced by external biotic cues such as the presence of roots or ectomycorrhizae (Fig 6). Both plant roots and fungal hyphae produce various exudates that chemoattract bacteria and can be used as a nutrient source (Johansson *et al*., 2004; Nazir *et al*., 2010; Stopnisek *et al*., 2016; Deveau *et al*., 2010) but bacteria differ in their abilities to use these nutrients (Frey *et al*., 1997). As suggest our results, in the absence of competition between bacteria, roots and ectomycorrhizae may support more bacterial biofilm formation than the fungus alone. However, competition between bacteria in natural environment is likely to hinder such behavior (Förster *et al*., 2016; Kastman *et al*., 2016) and it will be necessary to integrate the multiple interactions that occur within multispecies mixed biofilms in further studies. In addition, settings mimicking the physico-chemical properties of soils and their texture will allow to shed light onto the parameters that drive bacterial-fungal interactions in soils.

If there are evident advantages for bacteria to form biofilms (Balbontín *et al*., 2014; Stopnisek *et al*., 2016), the consequences of such biofilm formation for the fungi are not clear. On one hand, bacterial biofilms could protect the fungal hyphae against grazing, toxic compounds and buffer environmental variations (Frey-Klett *et al*., 2011; Kuramitsu *et al*., 2007). On the other hand, the presence of biofilms on the growing active area of fungal colonies may limit the capacity of fungi to degrade organic matter and induce a competition for nutrients between the bacterial community and the hyphae. In this second hypothesis, we expect that fungi, and particularly saprotrophic fungi, would have developed strategies to block the formation of biofilms on their hyphae (Stöckli *et al*., 2016; Essig *et al*., 2014). Our data support the hypothesis of de Carvalho and colleagues (de Carvalho *et al*., 2015), in which the mycelium of ECM fungi is rather permissive to bacterial biofilm formation although exceptions like *T. melanosporum* exist. Conversely, the inhibition of biofilm formation was more frequent among Ascomycota than Basidiomycota. Whether this result reflects a reality or is biased by the existence of species-specific interactions such as demonstrated for AM fungi will need to be further investigated (Toljander *et al*., 2006; Scheublin *et al*., 2010). The mechanisms behind the inhibition of biofilm formation by certain fungi remain also to be discovered. However, our data suggest that they rely on different processes depending on the fungal species. For instance, preliminary data in *T. melanosporum* suggest that the fungus would actively inhibit biofilm formation because biofilm formation was observed on dead hyphae (data not shown).

The large taxonomic distribution of bacteria able to form a biofilm on *L. bicolor* S238N hyphae suggest that there was a low degree of specificity of the bacteria towards the fungal host, and thus that the interaction potentially relied on a common mechanism. Surprisingly, we observed that all bacterial biofilms were structured by a network of filaments that seem to maintain cell together and to anchor biofilms to the surface of hyphae and of glass fibers (Fig 2, 4). These skeleton-like structures were highly reminiscent of the DNA extracellular traps produced by human neutrophils, plant roots and amoebae to capture bacteria (de Buhr *et al*., 2016; Zhang *et al*., 2016; Tran *et al*., 2016). Our data also strongly suggest that these skeletons would be made of eDNA. While the presence of eDNA in bacterial biofilms is now well demonstrated (Flemming and Wingender, 2010), it is often seen as an anamorphous material. Yet, such eDNA organization into filaments in bacterial biofilms have been sparsely reported (Böckelmann *et al*., 2006; Rose *et al*., 2015; Barnes *et al*., 2012; Tang *et al*., 2013; Gloag *et al*., 2013; Jurcisek and Bakaletz, 2007; Novotny *et al*., 2013; Tran *et al*., 2016; Liao *et al*., 2014). Together with our data, this suggests that eDNA based skeletons are a common feature of bacterial biofilms shared between Actinobacteria, Bacteroidetes, Firmicutes, α-, β- and γ- Proteobacteria.

The role of such eDNA filaments in the formation and functioning of bacterial biofilms remains elusive. Consistently with the hypothesis that eDNA would serve as a cohesive molecule that maintain bacterial cells together and anchor them to a surface (Das *et al*., 2010; Tang *et al*., 2013), the addition of DNAse at the inoculation time of the bacteria fully blocked the formation of biofilm (data not shown). Although eDNA skeletons may have additional functions in bacterial biofilms (Gloag *et al*., 2013; Doroshenko *et al*., 2014), it is noteworthy that a broad range of organisms belonging to the Animal, Plant, Protist and Eubacteria Kingdoms all uses DNA for additional purposes than coding genetic information. Thus DNA, thanks to its adhesive physico-chemical properties, may have an additional universal function that has been overlooked so far.

Overall, our work indicates that soil fungi, and most particularly ECM fungi, may often serve as a support to biofilm formation for a wide range of soil bacteria, in addition to be a source of nutrients and a potential vector for bacterial mobility. Such biofilm formation on fungal hyphae is likely to be modulated by numerous biotic factors including roots exudates and fungal activities. Besides the importance of eDNA based filaments for the building of these biofilms, the molecular dialog involved in the formation of biofilms on fungal hyphae and the nature of the interaction engaged between the microorganisms will need to be further investigated.

## Acknowledgements

This work was supported by the French National Research Agency through the Laboratory of Excellence ARBRE (ANR-11-LABX-0002-01) and by the Plant-Microbe Interfaces Scientific Focus Area in the Genomic Science Program, the Office of Biological and Environmental Research in the U.S. Department of Energy Office of Science. Oak Ridge National Laboratory is managed by UT-Battelle, LLC, for the U.S. Department of Energy under contract DE-AC05-00OR22725. We would like to thank Francis Martin for helpful discussions and comments on the manuscript. We thank Frédéric Guinet for helping with the preparation of *in vitro* mycorrhized seedlings, Anais Gilet for providing fungal strains, and Béatrice Palin for technical support along the experiments.

---

**Supplemental Fig. 1.** Experimental setup used to analyze biofilm formation during tripartite interaction between *P. fluorescens* BBc6, *L. bicolor* S238N and *Populus* tremula *x alba*. Mycorhizal seedlings were first produced using the *in vitro* sandwich co-culture system (Felten *et al*. 2009). One-month-old mycorrhizal seedlings were then used to analyze *in vitro* biofilm formation on ectomycorrhizae, non-mycorrhized roots, and extramatrical mycelium.

**Supplemental Fig. 2.** Biofilm formation of the hyphae of *L. bicolor* S238N by bacteria as visualized by confocal microscopy. Fungal hyphae were stained with Wheat Germ Agglutin-AlexaFluor 633 (red) and bacteria with DAPI (blue), except *P. fluorescens* BBc6, B. ginsengisoli E3BF7_7 and Dyella sp. E3BF9_7 that constitutively expressed GFP (green). eDNA filaments are highlighted by yellow arrows in caption boxes. Images were obtained via 2D maximum intensity projection of 3D confocal microscopy images Magnification 40x.

**Supplemental Fig. 3.** Elemental mapping of P (b, c), N (d, e) and Si (f, g) of BBc6 filaments and biofilm matrix on glass fibers obtained by EDS Spectrometry coupled to SEM imagery. Electron image: a, raw elemental map; b, d and f, overlay of electron image and elemental map : c, e, g

**Supplemental Fig. 4.** Biofilm formation by *P. fluorescens* BBc6-GFP on hyphae of soil fungi. Fungal hyphae were stained with Wheat Germ Agglutin-AlexaFluor 633 (red), bacterial cells were GFP-tagged and eDNA was stained with DAPI (blue). Due to poor staining of *T. terrestris*, *A. ustus and P. funiculosum* hyphae by Wheat Germ Agglutin-AlexaFluor 633, hyphae of these fungi were imaged using transmitted light, coloured in red. Images were obtained via 2D maximum intensity projection of 3D confocal microscopy images. Magnification 40x.

